# Commensalism in the *Mimiviridae* giant virus family

**DOI:** 10.1101/782383

**Authors:** Sandra Jeudy, Lionel Bertaux, Jean-Marie Alempic, Audrey Lartigue, Matthieu Legendre, Lucid Belmudes, Sébastien Santini, Nadège Philippe, Laure Beucher, Emanuele G. Biondi, Sissel Juul, Daniel J. Turner, Yohann Couté, Jean-Michel Claverie, Chantal Abergel

## Abstract

Acanthamoeba-infecting Mimiviridae belong to three clades: Mimiviruses (A), Moumouviruses (B) and Megaviruses (C). The uniquely complex mobilome of these giant viruses includes virophages and linear 7 kb-DNA molecules called “transpovirons”. We recently isolated a virophage (Zamilon vitis) and two transpovirons (ma_B_tv and mv_C_tv) respectively associated to B-clade and C-clade mimiviruses. We used the capacity of the Zamilon virophage to replicate both on B-clade and C-clade host viruses to investigate the three partite interaction network governing the propagation of transpovirons. We found that the virophage could transfer both types of transpovirons to B-clade and C-clade host viruses provided they were devoid of a resident transpoviron (permissive effect). If not, only the resident transpoviron was replicated and propagated within the virophage progeny (dominance effect). Although B- and C-clade viruses devoid of transpoviron could replicate both ma_B_tv and mv_C_tv, they did it with a lower efficiency across clades, suggesting an ongoing process of adaptive co-evolution. We performed proteomic comparisons of host viruses and virophage particles carrying or cleared of transpovirons in search of proteins involved in this adaptation process. These experiments revealed that transpoviron-encoded proteins are synthetized during the combined mimiviruses/virophage/transpoviron replication process and some of them are specifically incorporated into the virophage and giant virus particles together with the cognate transpoviron DNA. This study also highlights a unique example of intricate commensalism in the viral world, where the transpoviron uses the virophage to propagate from one host virus to another and where the Zamilon virophage and the transpoviron depend on their host giant virus to replicate, this without affecting the giant virus infectious cycle, at least in laboratory conditions.

**Author Summary:** The Mimiviridae are giant viruses with dsDNA genome up to 1.5 Mb. They build huge viral factories in the host cytoplasm in which the nuclear-like virus-encoded functions (transcription and replication) take place. They are themselves the target of infections by 20 kb-dsDNA virophages, replicating in the giant virus factories. They can also be found associated with 7kb-DNA episomes, dubbed transpovirons. We investigated the relationship between these three players by combining competition experiments involving the newly isolated Zamilon vitis virophage as a vehicle for transpovirons of different origins with proteomics analyses of virophage and giant virus particles. Our results suggest that relationship of the virophage, the transpoviron, and their host giant virus, extend the concept of commensalism to the viral world.

## Introduction

As of today, the *Mimiviridae* family appears composed of several distinct subfamilies (1–10) one of which, the proposed Megamimivirinae (7,10,11), corresponds to the family members specifically infecting Acanthamoeba (1,3–5,8). Members of this subfamilies, collectively refers to as “mimiviruses” throughout this article, can be infected by dsDNA satellite viruses called virophages only able to replicate using the already installed intracytoplasmic viral factory (6,12–15). These new type of satellite viruses constitute the Lavidaviridae family (16). In addition to virophages, the members of the mimiviruses can be found associated with a linear plasmid-like 7 kb-DNA called a transpoviron (13,16,17) making their mobilome uniquely complex among the known large DNA virus families.

Sputnik, the first discovered virophage, was found to infect the A-clade Mamavirus (12). Since it caused the formation of abnormal, less infectious, Mamavirus_A_ particles, it was initially proposed that virophages would in general protect host cells undergoing giant viruses’ infections. Such a protective role was quantitatively demonstrated for the protozoan *Cafeteria roenbergensis*, infected by the CroV (2)virus, in presence of the Mavirus virophage (18, 19). However, it was later recognized that some virophages replicated without visibly impairing the replication of their associated host virus. This is the case for the mimiviruses of the B- or C-clades when infected by the Zamilon virophage (15). On the other hand, members of the A-clade appeared non permissive to Zamilon replication (15). The resistance to Zamilon infection was linked to a specific locus proposed to encode the viral defence system MIMIVIRE (20–22) but the actual mechanisms governing the virulence of a given virophage *vis à vis* its host virus remain to be elucidated (21, 22). The rather ubiquitous transpovirons might also be involved in the protection of the mimiviruses against the deleterious effects of virophages infection. In this study we took advantage of a newly isolated virophage (Zamilon vitis) and of its ability to propagate different transpovirons to investigate the specificity of the transpoviron/host virus relationship. We also used B- and C-clades strains of mimiviruses originally devoid of transpoviron to investigate the possible role of the transpoviron in the context of virophage co-infections. Finally, we analyzed the proteome of virophage particles replicated on B- and C-clades host viruses, with and without resident transpoviron, to identify transpoviron proteins that could be involved in the co-evolution process allowing the transpovirons to be replicated by mimiviruses.

## Results

### Megavirus_C_ vitis, Moumouvirus_B_ australiensis, Moumouvirus_B_ maliensis and their mobilomes

Three samples recovered from various locations (France, Australia and Mali) produced lytic infections phenotypes when added to cultures of *Acanthamoeba castellanii* cells. Viral factories were recognizable in the amoeba cells after DAPI staining and we observed the accumulation of spherical particles visible by light microscopy in the culture medium (Fig. 1A). The corresponding virus populations were cloned and amplified as previously described (23). In all cases, negative staining electron microscopy (EM) images confirmed the presence of icosahedral virions of ∼450 nm in diameter with a stargate structure at one vertex, as was observed previously for all *Mimiviridae* infecting Acanthamoeba cells (3,15,24,25). For some clones of the Megavirus_C_ vitis giant virus, associated icosahedral virions of ∼70 nm diameter were also visible, suggesting the presence of virophages (Fig. 1B-F).

**Figure 1:**
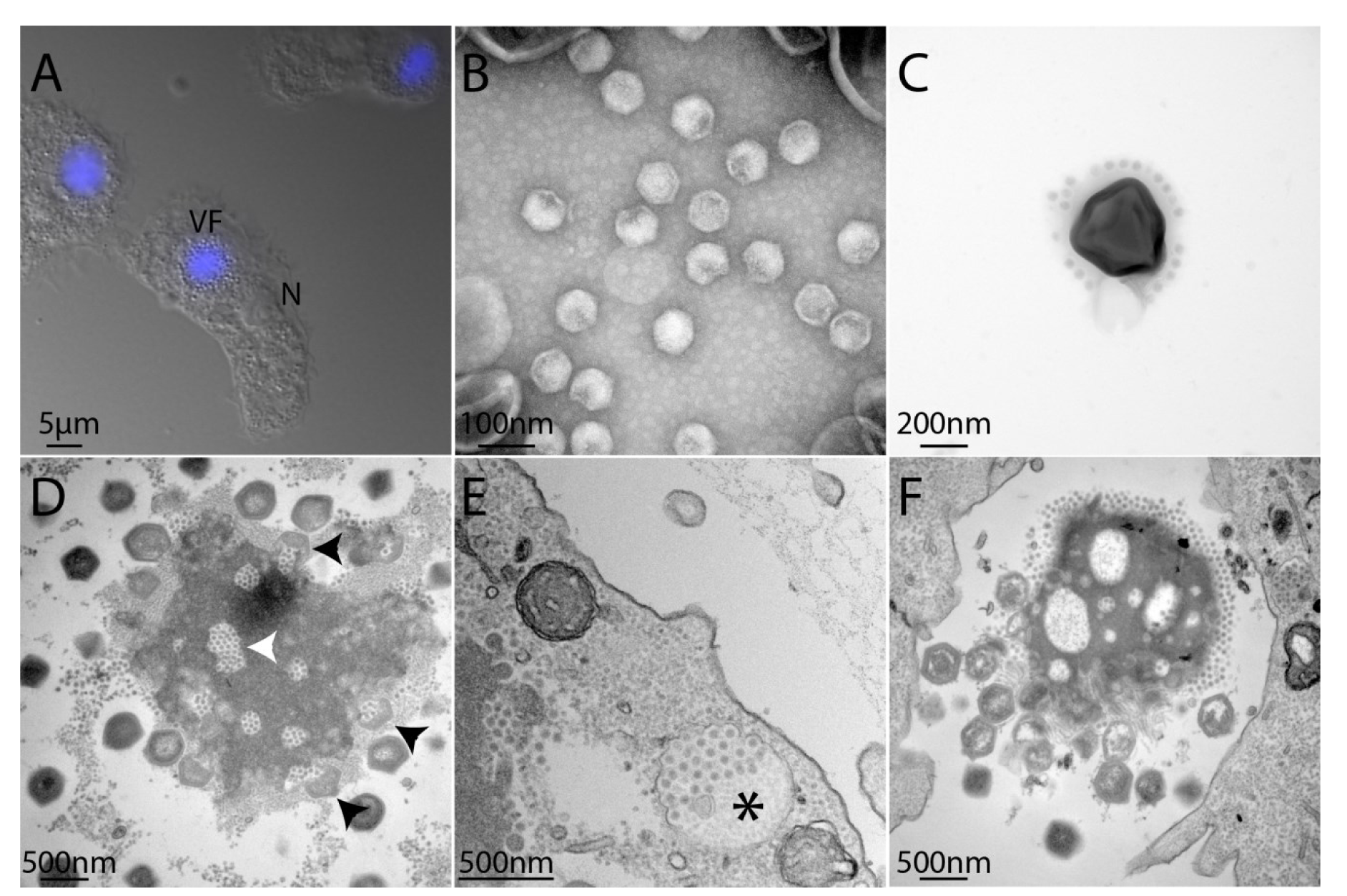
Microscopy images of Megavirus_C_ vitis and its associated virophage Z. vitis. (A) Fluorescence image of DAPI-stained *A. castellanii* cells infected by M_C_. vitis and its virophage. Viral particles are visible in the periphery of the viral factory (VF). The cell nucleus (N) remains visible but its fluorescence becomes undetectable due to the intense labeling of the VF DNA. (B) Transmission electron microscopy (TEM) Zamilon vitis particles observed by negative staining electron microscopy; (C) TEM of virophage particles stuck to the giant virus particle (negative staining); (D) Ultrathin section TEM of a M_C_. vitis viral factory observed in late infection of *A. castellanii* cells: virophage particles can be seen in holes in the VF (white arrowhead) as well as penetrating a maturing M_C_. vitis particle (black arrowheads); (E) Neo-synthesized Z. vitis virophage particles gathered in vacuoles (black star) are seen at the periphery of the infected cell suggesting that they are released by exocytosis. (F) TEM image of an isolated viral factory observed in an ultrathin section of a late infection of *A. castellanii* cells: virophages accumulate at one pole of the VF as well as in holes in the VF while immature and mature M_C_. vitis particles are seen at the opposite pole of the VF (Supplementary movie).

Based on its genome sequence, this new isolate was determined to belong to the C-clade and named Megavirus_C_ vitis. Its associated virophage was named Zamilon vitis, in reference to its genomic similarity with the original *Zamilon* virophage (15). In addition, the genome sequence assembly process revealed the presence of a 7kb dsDNA sequence homologous to previously described transpovirons (13). The Megavirus_C_ vitis associated transpoviron was named mv_C_tv. The Australian isolate was found to be a B-clade, and was named Moumouvirus_B_ australiensis. It came with its own transpoviron that we named ma_B_tv. Finally, the Bamako (Mali) isolate was determined to be another B-clade member, and named Moumouvirus_B_ maliensis. This last virus was devoid of transpoviron. Underscore A, B, and C are used throughout this article to indicate the clade origin of the various mimiviruses and of their natively-associated transpovirons.

### Comparison of Acanthamoeba cells co-infected by B- or C-clade mimiviruses and Z. vitis

The infectious cycles of M_C_. vitis, M_C_. chilensis, M_B_. australiensis and M_B_. maliensis appeared very similar to that of other mimiviruses, with an initial internalization of the virions in vacuoles, followed by the opening of the stargate and the fusion of the internal membrane with that of the vacuole to deliver the nucleoid in the cytoplasm. After 3h post infection (pi), viral factories develop in the cell cytoplasm, delineated by a mesh of fibres excluding all organelles (3, 26). Later on, neo-synthesized virions are seen budding and maturing at their periphery. Virophages specifically associated to the mimiviruses are thought to penetrate the cells at the same time as their host viruses, either enclosed in the host virus particles, or sticking to their external glycosylated fibrils (12) (Fig. 1 C). The virophages are devoid of a transcription machinery and thus use the transcription apparatus of the host virus to express their genome once released in the cell cytoplasm (12, 27). During infection of Acanthamoeba cells with M_C_. vitis or M_C_. chilensis in the presence of Z. vitis, regions depleted of electron dense material (“holes”) appeared in the viral factory prior to the assembly of any virion. Virophage particles then start to accumulate inside these holes (Fig. 1, Fig. S1 C-D, Supplementary movie) as early as 4h pi, before the production of host virus particles. Such infectious cycle is reminiscent of the one described for the association between virophage and mimiviruses (12,13,15), with virophages visible at the periphery of the viral factory, some of them seemingly penetrating inside the maturing giant virions (Fig. 1 D). The infectious cycle of Moumouvirus_B_ australiensis during coinfection of Acanthamoeba with the Z. vitis virophage was very similar to the one of M_C_. vitis except that instead of holes, the viral factory appeared to segregate the production of virophages in a separate compartment (Fig. S1 E-F). In all cases, during the latest stage, virophages were seen in large vacuoles that appeared to migrate toward the cell membrane to be released through exocytosis (Fig. 1 E). However, while Sputnik co-infections lead to aberrant and non-infectious host virus virions (12, 13), Z. vitis, as other Zamilon virophages, does not visibly impede the replication of its host viruses, abnormal particles of which were never observed (15).

### The newly isolated mimiviruses’ genomes

The dsDNA genome sequence of Megavirus_C_ vitis was assembled into a single contig of 1,242,360 bp with a G+C content of 25%. It was very close to Megavirus Terra1 (28) and to the C-clade prototype Megavirus_C_ chilensis (3) with whom it shared 99.1% and 96.9% identical nucleotides over the entire genome length, respectively (Fig. S2). Moumouvirus_B_ australiensis genome sequence was assembled in one contig of 1,098,002 bp (25% G+C), and that of Moumouvirus_B_ maliensis in one contig of 999,513 bp (25% G+C). As shown in Fig. S2, M_B_. australiensis and M_B_. maliensis belonged to B-clade and are closer to each other than to any other moumouviruses, thus initiating a third sub-lineage. Interestingly, the B-clade appeared the most divergent among mimiviruses.

### Zamilon vitis genome

In addition, we determined the 17,454 bp genome sequence (30% G+C) of Zamilon vitis. It was closely related to that of other virophages infecting the B- and C-clades, sharing 97.8% identical nucleotides with the prototype *zamilon* virophage (15). The 20 predicted proteins were all conserved in other virophages infecting mimiviruses, sharing 40 to 80% identical residues with Sputnik (12), their most distant homolog.

### The mv_C_tv and ma_B_tv transpovirons

Finally, we determined the genome sequences of the two new transpovirons. The M_C_. vitis transpoviron (mv_C_tv) DNA sequence was 7,417 bp (22% G+C) in length and closely related to the one associated with Megavirus_C_ courdo7 (98% identical nucleotides). The M_B_. australiensis transpoviron (ma_B_tv) DNA sequence was 7,584 bp in length (22% G+C) and was related to the one associated with Moumouvirus_B_ monve (89% identical nucleotides). The reconstructed phylogeny of all the known transpoviron genomes clearly showed that they fell into three distinct clusters, mirroring the tripartite clades structure of the host viruses from which they were isolated (Fig. 2).

**Figure 2:**
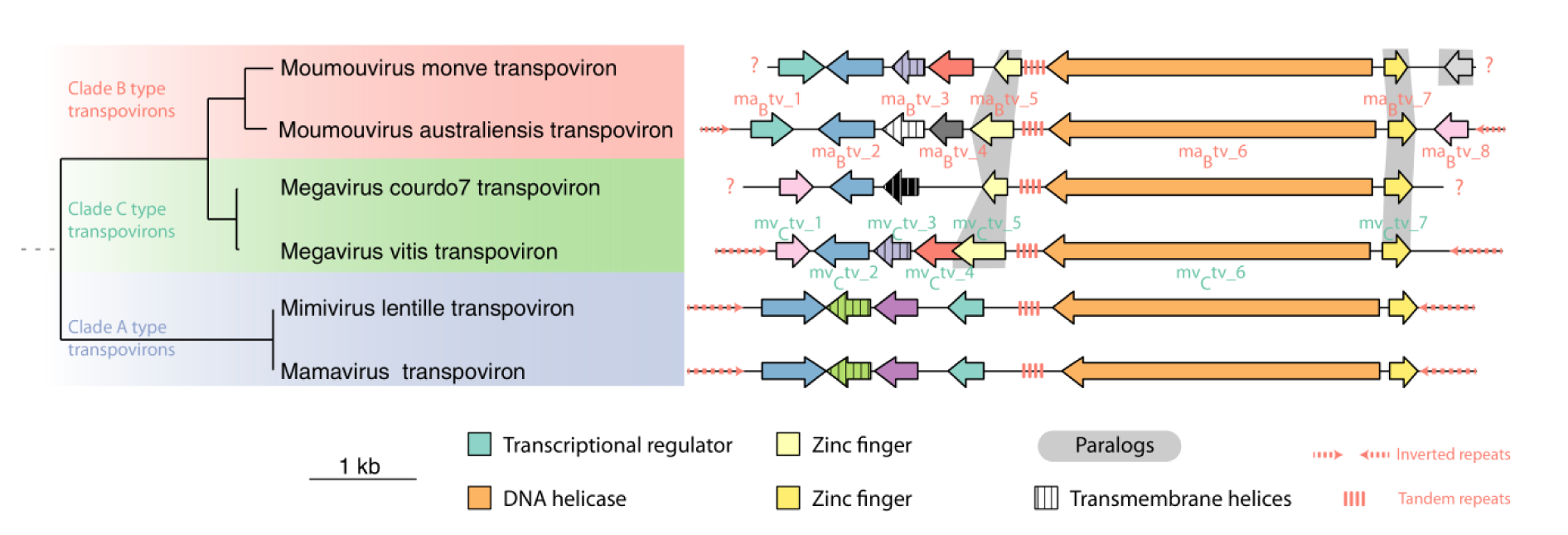
Phylogeny and genomic organization of transpoviron sequences. The phylogenetic tree (on the left) was computed from the concatenated sequences of shared orthologous predicted proteins using PhyML (29) with the LG+G model. Bootstrap values (not shown) are all equal to one. The genomic organization (right) shows orthologous genes represented with identical colors and paralogous genes (in a given genome) are highlighted in grey. Gene names are indicated for ma_B_tv and mv_C_tv.

Transpovirons exhibited terminal inverted repeats (TIRs) of 520 nt for Mamavirus_A_ and Lentille virus_A_ transpovirons, 380 nt for ma_B_tv and 510 nt for mv_C_tv. TIRs were missing from the transpovirons associated to Moumouvirus_B_ monve and Megavirus_C_ courdo7, most likely because their published sequences are incomplete (13). TIRs are well conserved within clades (90% of identical nucleotides between Mamavirus_A_ and Lentille virus_A_ transpovirons, lv_A_tv) but diverged between clades (56% identical nucleotides between lv_A_tv/mv_C_tv and 53% between mv_C_tv/ma_B_tv). A tandem repeat (TR) of 180-600 nt was present in the centre of all sequenced transpovirons, in an intergenic region 3’ from a conserved helicase (Fig. 2). These TRs were also well conserved within clades (80% identical nucleotides) and divergent between clades (39% identical nucleotides). It is worth mentioning that an evolutionary link between virophages and transpovirons has been proposed (30). Three predicted proteins were found in all transpovirons (*i.e.* core genes): a helicase (Mv_C_tv_6/Ma_B_tv_6), a protein of unknown function (Mv_C_tv_2/Ma_B_tv_2) and a zinc-finger domain containing protein (Mv_C_tv_7/Ma_B_tv_7), (Fig. 2). In addition, all transpovirons encode a small protein with a central transmembrane segment as their only recognizable similarity (Mv_C_tv_3/Ma_B_tv_3). Seven predicted proteins were shared by at least two transpovirons, including a transcriptional regulator conserved in clades A and B (Ma_B_tv_1), a protein paralogous to the above core zinc-finger-domain-containing protein and conserved in clades B and C (Mv_C_tv_5/Ma_B_tv_5), and five other predicted proteins without any functional signature. Five other predicted proteins were shared between at least two transpovirons but they do not bear any functional signature. Finally, four proteins of unknown function were unique and had no detectable homolog in the other transpovirons.

We analyzed the proteome of M_C_. vitis virions in search of transpoviron proteins specifically associated to this host virus. We identified three transpoviron proteins, Mv_C_tv_3, a putative membrane protein that could be anchored in the giant virus membrane, Mv_C_tv_2 and Mv_C_tv_4, two putative DNA-binding proteins (Fig. S3, Table S1).

**Table 1:** Permissivity of the host viruses to Z. vitis virophage and their selectivity for the transpovirons

| Clade | Giant virus | Permissivity to *Z. vitis* | Selectivity for transpoviron |  |
| --- | --- | --- | --- | --- |
|  |  |  | *Z. vitis* + *mv<sub>C</sub>tv* | *Z. vitis* + *ma<sub>B</sub>tv* |
| **A** | Mimivirus | - | *mv<sub>C</sub>tv*- | *ma<sub>B</sub>tv*- |
| **C** | *M<sub>C</sub>. chilensis* | + | ***mv<sub>C</sub>tv*+** | *ma<sub>B</sub>tv*+ |
|  | *M<sub>C</sub>. chilensis* + ***mv<sub>C</sub>tv*** | + | ***mv<sub>C</sub>tv*+** | ***mv<sub>C</sub>tv*+** *ma<sub>B</sub>tv*+ |
|  | *M<sub>C</sub>. chilensis* + *ma<sub>B</sub>tv* | + | *ma<sub>B</sub>tv* + ***mv<sub>C</sub>tv*+** | *ma<sub>B</sub>tv*+ |
| **C** | *M<sub>C</sub>. vitis* + <u>*mv<sub>C</sub>tv*</u> | + | ***mv<sub>C</sub>tv*+** | ***mv<sub>C</sub>tv*+** |
| **B** | *M<sub>B</sub>. australiensis* + <u>*ma<sub>B</sub>tv*</u> | + | ***ma<sub>B</sub>tv*+** | ***ma<sub>B</sub>tv*+** |
| **B** | *M<sub>B</sub>. maliensis* | + | *mv<sub>C</sub>tv*+ | ***ma<sub>B</sub>tv*+** |
|  | *M<sub>B</sub>. maliensis* + ***ma<sub>B</sub>tv*** | + | ***ma<sub>B</sub>tv*+** *mv<sub>C</sub>tv*+ | ***ma<sub>B</sub>tv*+** |
|  | *M<sub>B</sub>. maliensis* + *mv<sub>C</sub>tv* | + | *mv<sub>C</sub>tv*+ | *mv<sub>C</sub>tv* + ***ma<sub>B</sub>tv*+** |

*Transpovirons originally associated to a given host virus are underlined. The most represented transpovirons in the host population are shown in bold.*

Given that all known virophages infecting mimiviruses have been isolated in presence of a transpoviron, we also expected the presence of transpoviron-encoded proteins in Z. vitis virions. We thus analyzed the protein composition of the purified Z. vitis particles produced with M_C_. vitis. The proteomic study confirmed this prediction but suggest a specific interaction between the transpoviron proteins and the virophage in one hand, and the transpoviron proteins and the host virus particles in the other hand. Indeed, in virophage particles, contrary to what has been observed in M_C_ vitis particles, we consistently identified 2 different transpoviron proteins, Mv_C_tv_7, a putative DNA binding protein and in lesser amount the predicted helicase Mv_C_tv_6 (Table S1). Four additional proteins were also detected but in much smaller amount, three predicted DNA binding protein (Mv_C_tv_5, Mv_C_tv_2 and Mv_C_tv_4 in decreasing amounts) and a protein with unpredictable function, Mv_C_tv_1. These four proteins were also seen in the total proteome of the M_C_. vitis + Z. vitis virions. In contrast, they were all absent from the proteome of the cloned M_C_. Vitis particles (Fig. S3). Thus, the transpoviron encodes different subsets of proteins that might be specifically involved in their packaging in two alternative vehicles: the virophage or the host virus particle.

### Clade specificity of transpovirons

First, we verified that Z. vitis virophage replication was restricted to host viruses from the B- and C-clades, as previously described for Zamilon virophages (15) (Table 1). We also verified by PCR that the M_C_. vitis clone cleared from virophage and replicated on *A. castellanii* cells remained associated with its transpoviron mv_C_tv.

Purified virophage virions carrying the mv_C_tv transpoviron were then used to co-infect *A. castellanii* with two C-clade megaviruses (M_C_. vitis/mv_C_tv and M_C_. chilensis w/o transpoviron) and two B-clade moumouviruses (M_B_. australiensis/ma_B_tv and M_B_. maliensis w/o transpoviron) to assess whether the transpovirons were specific to a given clade of mimiviruses (Fig. 3).

**Figure 3:**
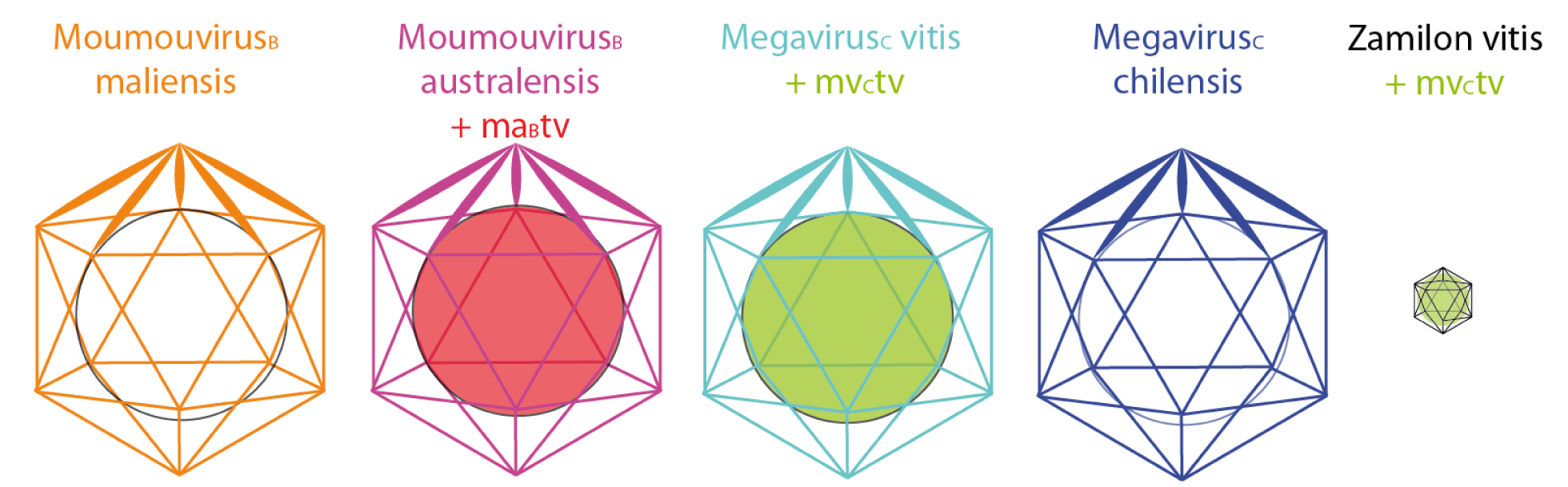
Viruses used in this study. M_B_. maliensis is represented in orange, M_B_. australiensis in purple, M_C_. vitis in cyan and M_C_. chilensis in dark blue. The transpovirons are represented as colored circles (green for mv_C_tv and pink for ma_B_tv) inside the giant virus and virophage capsids.

We performed specific PCR for each transpoviron after each round of co-infection and for each additional round of production (up to 10) to assess the presence of ma_B_tv and mv_C_tv DNA in the cultures after cell lysis. We also performed the proteomic analyses of the resulting virophages to assess the presence of transpoviron proteins (Table S1).

Co-infection of the virophage (carrying mv_C_tv) with M_B_. australiensis (carrying ma_B_tv) surprisingly produced a unique population of neo-synthesized virophage particles carrying the ma_B_tv transpovirons (Table 1, lane 6). The proteomic analysis of the purified virophage capsids also evidenced the replacement of the mv_C_tv proteins by their orthologues in ma_B_tv, ma_B_tv_7 and ma_B_tv_6 and to a lesser extend ma_B_tv_2. In contrast the orthologues of Mv_B_tv_1 (Ma_B_tv_8) and Mv_C_tv_5 (Ma_B_tv_5) were not detected (Table S1). Moreover, PCR performed along the infection cycle of M_B_. australiensis (carrying ma_B_tv) and the virophage (carrying mv_C_tv) did not show an increase in mv_C_tv while the ma_B_tv genome was clearly replicated (Fig S4A). These results suggested that the host virus strongly favour the replication of its natively associated transpoviron (Table I, Fig. 4A). If a different one is brought in by the virophage, it is lost and replaced by the one replicated by the host virus, a result we refer to as the “dominance effect”. Consequently, two populations of virophages carrying either mv_C_tv or ma_B_tv were at our disposal. We confirmed the dominance effect by co-infection of M_C_. vitis carrying mv_C_tv with virophages carrying ma_B_tv (Table 1, lane 5). Again, we observed the replacement in the virophage particles of ma_B_tv (DNA and proteins) by mv_C_tv (DNA and proteins). We also confirmed that the mv_C_tv genome was actively replicated while the amount of ma_B_tv genome remained stable along the infectious cycle (Fig S4B). We then used the virophages carrying either mv_C_tv or ma_B_tv to infect transpoviron-free B-clade (M_B_. maliensis) and C-clade (M_C_. chilensis) host viruses. We found that the virophage succeeded in transmitting each transpoviron to each “empty” B- or C-clade host viruses (Table 1, lane 2 and 7), a result we refer to as the “permissive effect”. However, we observed that ma_B_tv was preferentially replicated by M_B_. maliensis and mv_C_tv by M_C_. chilensis (Table 1). By cloning we showed that the resulting populations of B- and C-clade host viruses were mixtures of transpoviron positive and transpoviron negative particles. Furthermore, virophage particles produced by transpoviron negatives clones were also devoid of transpoviron (DNA and proteins), indicating that although transpovirons can be carried by virophages, their replication is performed and controlled by the host virus.

**Figure 4:**
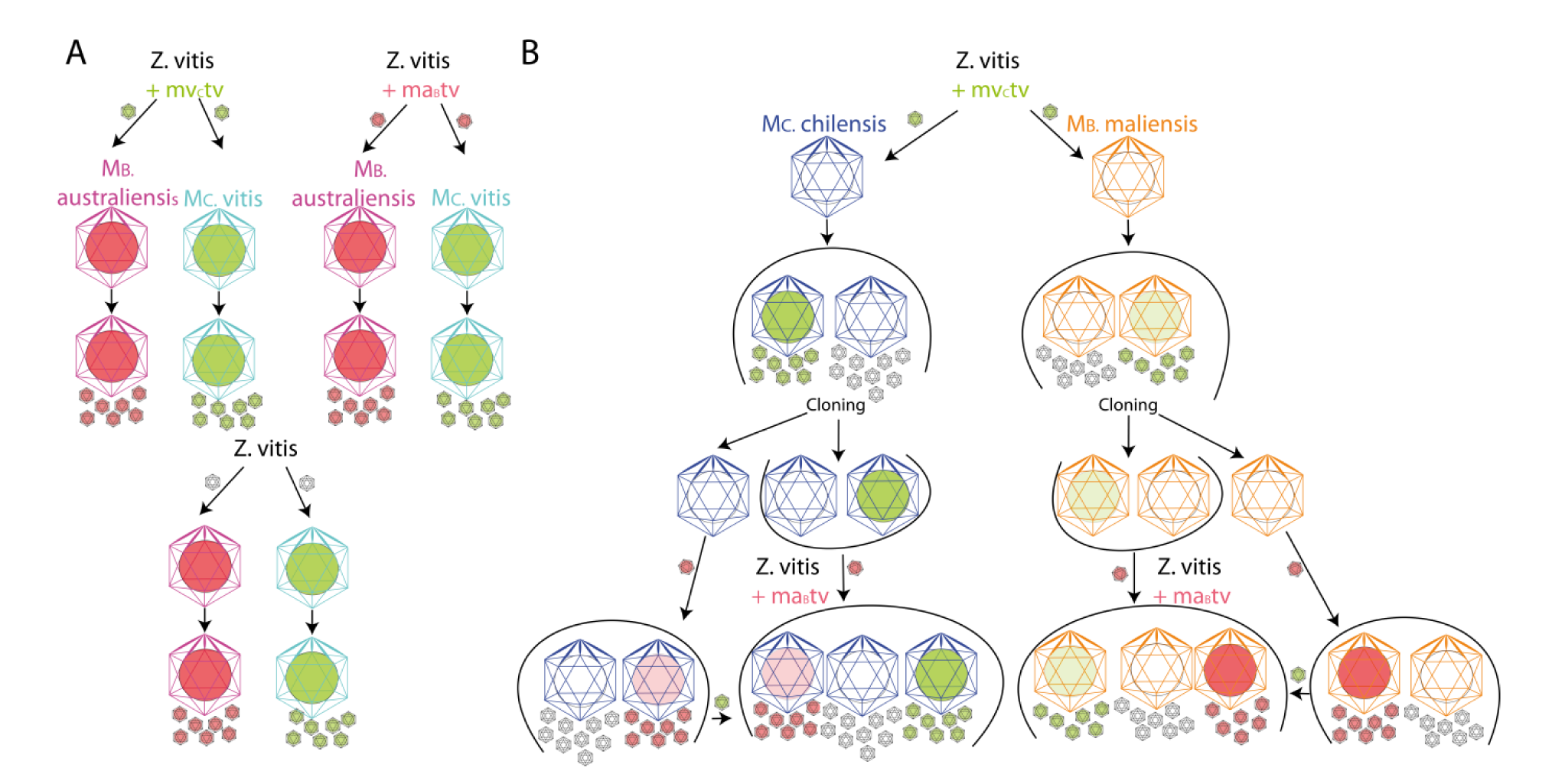
B- or C-clade host viruses co-infected with virophages carrying ma_B_tv or mv_C_tv transpovirons A] **Dominance effect** of the resident transpoviron (mv_C_tv in M_C_. vitis, green circle in cyan capsid; ma_B_tv in M_B_. australiensis, pink circle in purple capsid) over the one carried by Z. vitis. Empty Z. vitis (black contour, white capsid) do acquire the resident transpoviron upon replication; B] **Permissive effect**: the two type of transpovirons can be imported and replicated by empty M_C_. chilensis (dark blue capsid, white circle) and M_B_. maliensis (orange capsids, white circle), although with different efficiencies. Color intensities of the circles (pink for ma_B_tv, green for mv_C_tv) illustrate the abundance of the transpovirons in the host and virophage particles.

We finally took advantage of the permissive effect to produce two populations of clade B (M_B_. maliensis) and C (M_C_. chilensis) host viruses, each carrying the ma_B_tv or mv_C_tv transpoviron. We then challenged them using virophages carrying the other transpoviron. We performed PCR specific for either transpoviron after each round of virophage co-infection (up to 10 successive rounds of virophage production) and assessed their presence after cell lysis.

The proteome of the virophages particles was also analysed in order to assess the presence of transpoviron proteins possibly associated to the transpoviron DNA. The results of the various experiments are presented in Table 1 and Table S1 and are interpreted in Figure 4.

When we infected populations of M_C_. chilensis carrying ma_B_tv or mv_C_tv with virophages carrying the complementary transpoviron, the resulting population of M_C_. chilensis and virophages became positive for the two transpovirons (Table 1, lane 3 and 4). This apparently violated the strict “dominance effect” observed with M_C_. vitis and M_B_. australiensis. However, the persistence of a subpopulation of empty M_C_. chilensis virion (i.e. devoid of transpoviron) might also explain our results without refuting the dominance rule. In that case, the replication and propagation of the competing virophage-borne transpoviron would be performed by the transpoviron-null (empty) M_C_. chilensis subpopulation. We investigated this possibility by cloning virions from the mixed mv_C_tv/ma_B_tv M_C_. chilensis population and examining their transpoviron content. Strikingly, we never observed M_C_. chilensis virions simultaneously carrying both types of transpovirons. On the other hand, we observed either mv_C_tv positive (mv_C_tv+) or ma_B_tv positive (ma_B_tv+) M_C_. chilensis clones, as well as others devoid of transpovirons (Fig. 4). This result also suggests that the association between the host virus and the transpovirons are not stable when resulting from a first encounter. In the case of M_B_. maliensis in the presence of virophage ma_B_tv, the replacement of the mv_C_tv transpoviron from host particles appeared to be faster, which resulted in the rapid disappearance of mv_C_tv after only 6 rounds of replication. We also observed that the loss of the transpoviron over host virus replication, in the absence of virophage, was more rapid for M_B_. maliensis than M_C_. chilensis suggesting the association between the host virus and the transpoviron was not stable. The cloning step provided virophage-free host virus clones with which to replicate these competition experiments: the ma_B_tv+ or mv_C_tv+ clones infected by virophages carrying the complementary transpoviron again produced a mixed population of ma_B_tv+, mv_C_tv+ and transpoviron-null virions (Table 1, lane 8 and 9). Thus, the persistence of particles devoid of transpoviron allows us to conclude that our results are a combination of the “dominance effect” applied to the subpopulation of transpoviron positive virions and of the “permissive effect” applied to the transpoviron-null subpopulation. In the resulting virophage particles, the only transpoviron proteins consistently identified were MvCtv_7/MaBtv_7 and MvCtv_6/MaBtv_6. As expected, they were absent from virophages devoid of transpoviron (Table S1).

To elucidate whether the transpoviron could have a protective role against infection of the host virus by the virophage, we compared the infectious cycles of Acanthamoeba cells infected by M_C_. chilensis carrying ma_B_tv, mv_C_tv or without transpoviron. They were strikingly similar both in terms of cycle length and virus production yields. Transpovirons are thus not key, at least in laboratory conditions, in regulating the permissivity of mimiviruses to virophage infection.

## Discussion

Mimiviruses are unique in their association with two distinct (often co-existing) dependent entities, virophages and transpovirons, somewhat reminiscent of phages and plasmids afflicting bacteria. As for the virophage, the presence of host virus-like regulatory elements (terminator hairpin, late promoter (31–34)) flanking the transpoviron genes suggest that they also use the host virus transcription machinery rather than that of the cell. The transpoviron might also rely on the host virus DNA replication machinery, in absence of transpoviron-encoded DNA polymerase. Our competition experiments between the mv_C_tv *versus* ma_B_tv transpovirons resulted in the replication of only one transpoviron. Interestingly, the “winner” corresponds to the type originally associated to the host virus (mv_C_tv for M_C_. vitis and ma_B_tv for M_B_. australiensis, Table 1), a phenomenon we called the “dominance effect”. This finding was also confirmed by the immediate replacement of mv_C_tv by ma_B_tv proteins in virophage particles synthetized with M_B_. australiensis. However, this result is not simply due to a strict clade-wise specificity. The use of transpoviron-free host virus particles allowed us to demonstrate that M_C_. chilensis and M_B_. maliensis can replicate and incorporate each transpoviron, independently (Table 1). Yet, we observed a marked difference in permissivity, with B- and C-clade host viruses favouring their cognate transpoviron types. The central TRs sequences and the TIR flanking the transpovirons replicated by A-clade *versus* B- or C-clade host viruses are markedly different. These differences might cause the lack of replication of ma_B_tv and mv_C_tv in the A-clade Mimivirus. The lesser differences between the B- and C-clade transpovirons might then explain why both of them can still be replicated by M_B_. maliensis and M_C_. chilensis. In these host viruses, the competition experiment resulted in the simultaneous replication of both transpovirons. However, the sub-cloning of the mv_C_tv+/ma_B_tv+ population resulted in mv_C_tv+ *only*, ma_B_tv+ *only*, or transpoviron negative clones. It also appears that neither of the two transpovirons remains stably associated with a host virus for which it was a first encounter, while a preference could emerge once a stable association has been established by coevolution (*i.e.* M_B_. australiensis with ma_B_tv, M_C_. vitis with mv_C_tv). Finally, in virophage particles there was fewer copies of ma_B_tv either produced with M_B_. australiensis, M_B_. maliensis or M_C_. chilensis and fewer copies of mv_C_tv produced with M_C_. chilensis or M_B_. maliensis, compared to the number of copies of mv_C_tv produced with M_C_. vitis. Different transpovirons, even efficiently replicated by the host virus, thus appear to be loaded at different efficiencies in the virophage particles (Fig. S5). A similar result was previously described for the Lentille virus transpoviron that could only be detected in Sputnik2 virophage particles using FISH experiments (13). A consequence of such suboptimal associations was the production of virophages devoid of transpovirons that could then be used to identify candidate proteins involved in the transpoviron/virophage association. The only difference between virophage particles carrying or not transpovirons was the recurrent presence of two transpoviron-encoded proteins (Mv_C_tv_7/Ma_B_tv_7 and Mv_C_tv_6/Ma_B_tv_6) together with the DNA molecule as an episome (Table S1, Fig. S5) suggesting the virophage was a mere vehicle for the transpoviron. These proteins are conserved in all transpovirons, are predicted to be DNA-binding, and were not identified in the proteome of the host virus. Instead, the most abundant transpoviron proteins in M_C_. vitis virions were two predicted DNA-binding proteins (Mv_C_tv_2, Mv_C_tv_4) and one predicted membrane protein (Mv_C_tv_3) that could be anchored in the host virus membrane (Table S1). The sequence of this short protein is not conserved between transpovirons associated to different host virus clades but all transpovirons encode such predicted membrane protein. The dominance effect is reminiscent of plasmids incompatibility or entry exclusion for related plasmids (35–38) or superinfection immunity (39) and superinfection exclusion (40) used by prophages to prevent superinfection by other bacteriophages. For the latter, DNA-injection is inhibited by a membrane-bound protein (41–43), paralleling the eventual role of the transpoviron-encoded membrane protein in giant virus particles carrying a transpoviron.

Since the transpoviron genome is present in both the host virus and the virophage particles, the transpoviron DNA might also adopt different organization depending on the vehicle (host virus or virophage particles) used for its propagation. The Mv_C_tv_7/mMa_B_tv_7 and Mv_C_tv_6/Ma_B_tv_6 proteins could be involved in the packaging or delivery of the transpoviron in and from the virophage particle, while the Mv_C_tv_2/Ma_B_tv2 and Mv_C_tv_4 proteins could play a similar role *vis à vis* the host virus and the Mv_C_tv_3 membrane protein may play a role in the dominance effect. Further studies are needed to elucidate the mechanisms at work in packaging and delivery of transpoviron genomes as well as in transpoviron dominance.

The first two types of virophage that have been discovered, Sputnik and Mavirus, respectively infecting Mimivirus (12) and Cafeteria roenbergensis (18) are strongly deleterious to their host viruses, diminishing the production of infectious particles (12,13,44) or stopping it altogether (18, 19), effectively protecting the cellular hosts. As parasite of another parasite (the giant virus), these virophages are *bona fide* hyperparasites (45, 46). The detection of many additional virophage-related sequences in aquatic environment together with that of mimivirus-like viruses suggested that they might have a significant ecological role in regulating the population of the giant viruses and of their cellular host (micro-algae or heterotrophic protozoans) (47–49). However, the hypovirulence (of the host virus) induced by the hyperparasite may ultimately limit its own reproductive success (45). Thus the evolutionary trajectory of the virophage/host virus/cellular host parasitic cascade may remain antagonistic or end up in a mutualistic or commensal relationship. The uniquely complex tripartite parasitic cascade transpoviron/virophage/host virus analyzed in this work, where none of the actors appears to have a detrimental effect on the others, at least in laboratory conditions, might be at a neutral equilibrium reached as a stable compromise after eons of intricate antagonistic evolution. To the best of our knowledge, the relationship between the transpoviron, the Zamilon virophage, and their host giant virus analyzed in this work represents the first example of bipartite commensalism in the viral world.

## Materials and Methods

### Viruses isolation

The new giant viruses’ strains were isolated from soil recovered from a French vineyard (43°20’25”N-5°24’51”E, M_C_. vitis), muddy water collected in the Ross river mangrove near Townsville (19°15’42.39“S, 146°49’5.50”E, M_B_. australiensis), and water collected from a well nearby Bamako (12°31’28.6“N 7°51’26.3”W, M_B_.maliensis). After being treated with antibiotics the different resuspended samples were added to a culture of *Acanthamoeba castellanii* (Douglas) Neff (ATCC 30010^TM^) cells adapted to Fungizone (2.5 μg/mL) and cultures were monitored for cell death as previously described (3,23,50,51). Each virus was then cloned (23) and the viral clones (M_B_. australiensis, M_C_. vitis and M_B_. maliensis) were recovered and amplified prior purification, DNA extraction and cell cycle characterization by electron microscopy. Permissivity to virophage infection on the various giant viruses were analyzed using the same protocol to assess the production of virophage after one round of co-infection of each clone with the purified virophage.

### Virus purification

All giant viruses were purified on a CsCl gradient as previously described (3). In contrast to Sputnik virophage, Z. vitis and our various giant viruses were still infectious after treatment at 65°C. We thus could not apply the previously published protocol (13) to separate Z. vitis from the giant viruses. Instead we used several steps of filtration/centrifugation with a final purification on sucrose gradient. The preparation was filtered on 0.2 µm filter and centrifuged at 100,000 g for 90 minutes. The pellet was resuspended in 5 mL of 40 mM Tris HCl pH7.5 and filtered on 0.1 µm filter. The filtrate was then centrifuged at 150,000 g for 1h, and the pellet resuspended in 0.2 mL of 40 mM Tris HCl pH 7.5 and loaded on a 70%/60%/50%/40% sucrose gradient and centrifuged at 200,000 g for 24h. The band corresponding to the virus was recovered with a syringe and washed once with 40 mM Tris HCl pH 7.5. The purification was controlled by negative staining observation using a FEI Tecnai G2 operating at 200 kV (Fig. S6). Competition experiments were performed using a large excess of virophage particles compared to the giant virus (10^3^ for 1).

### Synchronous infections for TEM observations of the infectious cycles

*A. castellanii* adherent cells in 20 mL culture medium were infected with each giant virus with a MOI of 50 for synchronization. After 1h of infection at 32°C, cells were washed 3 times with 30 mL of PPYG to eliminate the excess of viruses. For each infection time (every hour from 1h to 11h pi), 2.5 mL were recovered and we did include them in resin using the OTO (osmium-thiocarbohydrazide-osmium) method (52). Ultrathin sections of 90 nm were observed using a FEI Tecnai G2 operating at 200 kV.

### DNA extraction for sequencing

For the M_C_. vitis clone still associated with the Zamilon vitis virophage we did not try to exhaustively separate the giant virions from Z. vitis virions prior to DNA extraction. For each giant virus clone, genomic DNA was extracted from 10^10^ virus using the PureLink^TM^ Genomic DNA mini kit (Invitrogen) according to the manufacturer’s protocol. Finally, for the virophage, after its separation from the giant virus, the DNA was extracted using the same protocol. The purified DNAs were loaded on an agarose gel in search of an extra DNA band suggestive of the presence of an episome which could correspond to a transpoviron.

The M_B_. maliensis purified DNA was sent to the Novogene Company for library preparation and Illumina PE150 sequencing.

#### Library preparation for Nanopore technology

The ma_B_tv transpoviron DNA was extracted and purified from an agarose gel (Supplementary methods). DNA was quantified using a Qubit 3.0 Fluorimeter (Thermo Fisher Scientific, MA, USA) following the manufacturer’s protocol, and was found to be 12.7 ng.mL^-1^. 7.5 µL of this DNA was used for library preparation using the RAD002 kit (Oxford Nanopore Technologies, Oxford, UK). Since the input quantity of DNA was lower than recommended for this kit, the active FRM reagent was diluted with three volumes of heat-inactived FRM, to avoid over-fragmentation of the DNA. The library preparation reaction was set up as follows: The reaction (DNA 7.5 µL, 0.25 x FRM 2.5 µL) was incubated for 1 minute at 30°C followed by 1 minute at 75°C. We added 1 mL of RAD reagent from the RAD002 kit and 0.2mL of Blunt TA ligase (New England Biolabs, MA, USA) and the reaction was incubated for 5 minutes at room temperature. The prepared library was then loaded onto a FLO-MIN106 flowcell (version 9.4 nanopores) as per Oxford Nanopore Technologies’ standard protocol.

#### Library preparation for Illumina technology

Genome sequencing was performed using the instrument Illumina MiSeq. Libraries of genomic DNA were prepared using the Nextera DNA Library Preparation Kit as recommended by the manufacturer (Illumina). The sequencing reaction was performed using the MiSeq Reagent Kit v2 (300-cycles), paired end reads of 150nt x 2 (Illumina).

### Genome sequencing, assembly and annotation

The assembly of Moumouvirus_B_ australiensis genome sequence was performed on one SMRT cell of pacbio data using the HGAP workflow (53) from the SMRT analysis framework version 2.3.0 with default parameters, resulting in 84,565 corrected reads.

The MinION library was sequenced for 1 hour on a MinION Mk1B flowcell (Oxford Nanopore Technologies), generating approximately 220 Mb of sequence data. Basecalling was performed using the *1D Basecalling for FLO-MIN106 450bps r1.121* [*Workflow ID: 1200*] workflow (Oxford Nanopore Technologies), and yielded 128,623 reads with a mean length of 1,701 bases. Data was filtered to remove reads with a quality score below 8, leaving 76,936 reads, and a mean length of 2,369 bases. The reads were assembled using Canu with the default parameters, but with the option stopOnReadQuality=false. The Moumouvirus_B_ australiensis transpoviron sequence (ma_B_tv) resulting from this assembly was further polished using the M_B_. australiensis pacbio error corrected reads.

The assembly of the Megavirus_C_ vitis, Zamilon vitis and Megavirus vitis transpoviron genomes (mv_C_tv) was performed using Spades (54) (version 3.9.0) on MiSeq Illumina paired-end reads and Pacbio long reads when available. Spades was used with the following parameters: careful, k=17, 27, 37, 47, 57, 67, 77, 87, 97, 107, 117, 127, cov-cutoff=off and pacbio option. For Moumouvirus_B_ maliensis, for which long reads were not available, the assembly was performed using Illumina paired-end reads and Spades (version 3.12.0) with the following parameters: careful, No reads correction, k=33,55,77,99,127.

Gene annotation of genomic sequences was done using Augustus (55) trained on already published members of the subfamily gene sets. The tRNAs were searched using tRNAscan-SE (56) with default parameters. Functional annotation of predicted protein-coding genes was done using homology-based sequence searches (BlastP against the NR database (57) and search for conserved domains using the Batch CD-Search tool (58)).

### Phylogenetic analyses

The phylogenetic tree of the transpovirons (Fig. 2) was computed using a concatenation of the conserved genes. The optimal model “LG+G” was selected using Prottest (59). Phylogeny of the giant viruses (including Megavirus_C_ vitis, Moumouvirus_B_ australiensis, Moumouvirus_B_ maliensis, Fig. S2) was performed on a concatenation of genes shared by all mimiviruses as predicted by CompareM (https://github.com/dparks1134/CompareM). After protein clustering using MCL (60), clusters containing exactly one gene per virus were aligned using Mcoffee (61) and concatenated. We next produced a phylogenetic tree using Prottest (59) and the resulting superalignment. The “VT” model was considered the optimal model. Average nucleotide identity between the different strains (Fig. S2) was calculated using the OAU tool (62).

### MS-based proteomic analyses

#### Characterization of virion proteomes by data dependent acquisition

For proteomic analysis the virions pellets were resuspended in Tris HCl 40 mM pH 7.5, 60 mM DTT, 2% SDS and incubated 10 min at 95°C. The protein concentration of the lysates were measured using the 660 nm Protein Assay Reagent appended with Ionic Detergent Compatibility Reagent (Thermo Scientific) and 4 µg of proteins were analyzed as previously described (23). Briefly, proteins were stacked in the top of a 4–12% NuPAGE gel (Invitrogen) before R-250 Coomassie blue staining. Proteins were then in-gel digested using trypsin (sequencing grade, Promega). Resulting peptides were analyzed by online nanoLC-MS/MS (Ultimate 3000 RSLCnano and Q-Exactive Plus or HF) using a 120-min gradient. Two replicates were analyzed per sample, except for Z. vitis purified from M_B_. maliensis and Z. vitis purified from M_C_. chilensis containing ma_B_tv. Peptides and proteins were identified and quantified using MaxQuant software (63) (version1.6.2.10). Spectra were searched against the corresponding Megavirus_C_ chilensis, Megavirus_C_ vitis, Moumouvirus_B_ australiensis, Moumouvirus_B_ maliensis, Zamilon vitis, mv_C_tv, ma_B_tv and *Acanthamoeba castellanii* protein sequence databases and the frequently observed contaminants database embedded in MaxQuant. Minimum peptide length was set to 7 aa. Maximum false discovery rates were set to 0.01 at PSM, peptide and protein levels. Intensity-based absolute quantification (iBAQ) (64) values were calculated from MS intensities of unique+razor peptides. Proteins identified in the reverse and potential contaminants databases as well as proteins only identified by site were discarded form the final list of identified proteins. For each analysis, iBAQ values were normalized by the sum of iBAQ values in the analyzed sample (65). Only proteins identified with a minimum of 2 unique+razor peptides in one sample were considered.

### Dominance effect validation by PCR

Cells were grown in T75 flasks and infected either with M. australiensis (carrying ma_B_tv) at MOI 10 and a large excess (10^3^ for 1 giant virus particle) of virophage carrying mv_C_tv, or with M. vitis (carrying mv_C_tv) at MOI 10 and a large excess of virophage carrying ma_B_tv. After 40 min of infection, the cells were washed three times and distributed in 12-well plates (1mL per well). Cells were collected from a well at 45min, 2h, 4h, 6h, 8h and 24h post-infection. The cells were centrifuged at 1,000 x g for 3 min, except for the last point at 24h which was centrifuged at 16,000 x g for 10 min. Each pellet was resuspended in 100 µL of PPYG medium and cells were frozen in liquid nitrogen to stop the infection and stored at −80°C.

PCR were performed using the *Terra* PCR Direct *Polymerase* Mix (Clontech) directly on the cell lysates using the following primers:

qPCR-matv_F TCGCTCATTGATTCACTTTGTAC; qPCR-matv_R AATGTATTATGGGCGAATAATGTT; PCR produced an amplicon of 185bp

qPCR-mvtv_F GGCATAAGCAGGTTCGAAAT, qPCR-mvtv_R CATGGCGTGATATTGGTGTG; PCR produced an amplicon of 194bp

The PCR experiments were stopped after 20 cycles of amplification and 7µL of the reaction products were deposited on agarose gel (Fig. S4)

### Competition experiments

Cells were grown in T25 flasks (5 mL growth medium) and infected with host viruses carrying one transpoviron) at MOI 0.25 and a large excess (10^3^ for 1 giant virus particle) of virophage carrying the complementary transpoviron. After cell lysis, 100 µL of the culture medium containing virophage and host viruses were used to infect another T25 flask containing adherent fresh cells. This process was repeated 10 times.

### Selective identification of transpoviron in virions capsids

To distinguish the mv_C_tv and ma_B_tv transpovirons, two sets of primers were designed:

mv_C_tv_PFwd_: ACCTTCTTGTGCCTTTACTGC, mv_C_tv_PRev_: CAGGGTTCGGACGGATTACT; PCR produced a 939 bp amplicon.

ma_B_tvP_Fwd_: TCGCTCATTGATTCACTTTGTAC, ma_B_tvP_Rec_: CAAAGGGGAGGAAATAATGGAGA; PCR produced a 263 bp amplicon.

PCR were performed using the *Terra* PCR Direct *Polymerase* Mix (Clontech) directly on the cell lysates after each round of co-infection. To check the stability of the transpovirons over time, up to 10 additional rounds of virus production were performed and the presence of the transpoviron was assessed by PCR.

Total DNA was extracted from purified host viruses and virophages using the PureLink Genomic DNA Mini Kit (ThermoFischer Scientific) and serial dilutions of DNA were performed and deposited on a 0.8 % agarose gel ran for 45 min at 100V. For host viruses, we deposited from 1 to 0.25 µg of total DNA and for virophages, from 1 µg to 62.5 ng DNA. DNA bands were revealed using BET staining and images were recorded on a Chemi-smart 2000WL-20M camera (Fischer Bioblock Scientific).

## Supporting information captions

**Figure S1: TEM comparison of viral factories of M_C_. vitis alone (A-B) with M_C_. vitis (C-D) or Moumouvirus_B_ australiensis (E-F) co-infected by Z. vitis.** (A-C) Early VF of M_C_. vitis and M_C_. vitis/Z. vitis coinfection prior giant virus virions production. (C) Holes are clearly visible in the VF when Z. vitis is infecting M. vitis and some virophages can be seen (inset) at the periphery of the VF. (B-D) Late VFs. M_C_. vitis virions are clearly visible at the periphery of the VFs and Z. vitis particles can be seen in some holes (inset) (D). (E-F) Late M_B_. australiensis VFs producing giant virus particles and virophages at two different poles of the VF.

**Figure S2: Genomic sequence conservation and phylogeny of the mimiviruses.**Phylogenetic tree (on the left) was computed from the concatenation of shared orthologous genes peptide sequences using PhyML (1) with the VT model. The heatmap shows the average nucleotide sequence identity between the mimiviruses fully sequenced genomes estimated using the OrthoANIu tool (2).

**Figure S3: Proteome comparison of Mv_C_. Vitis purified capsids** resulting from infection of Acanthamoeba cells in the presence or absence of Z. vitis. For the latter, the virophage capsids were not separated from the ones of the giant virus. The abundances of M_C_. vitis (blue), Z. vitis (orange) and mv_C_tv (purple) proteins were compared based on their extracted iBAQ values normalized on the most abundant M_C_. vitis protein (Mvi_626). Higher values correspond to lower protein abundances.

**Figure S4: Dominance effect:** PCR specific of each transpoviron were performed using ma_B_tv and mv_C_tv specific primers *A. castellanii* cell lysates along the infectious cycle of (A) M_B_. australiensis/ma_B_tv co-infected with a virophage carrying mv_C_tv; (B) M_C_. vitis/mv_C_tv co-infected with a virophage carrying ma_B_tv. The transpoviron brought is in a large excess at the beginning of the infection (can be detected in the very early time points). The host virus is clearly replicating the transpoviron it carries after 2h pi, while the one brought in by the virophage does not appear to be replicated.

**Figure S5: Agarose gel of DNA extracted from purified virophages.** Serial dilutions of DNA extracted from purified virophages associated with ma_B_tv or mv_C_tv and produced with empty M_C_. chilensis (left gel), M_B_. maliensis (right gel) and M_B_. australiensis/ma_B_tv or M_C_. vitis/mv_C_tv (middle gel). The mv_C_tv transpoviron is clearly more efficiently associated with the virophage particles than ma_B_tv, whether produced with M_C_. vitis, M_C_. chilensis or M_B_. australiensis host viruses.

**Figure S6: Virophage purification on sucrose gradient and negative staining images of the recovered disk.** Negative staining confirms that the white disk corresponds to debris (A) and the blue disk to the purified virophage (B).

**Supplementary Movie: 3D reconstruction of an Acanthamoeba cell co-infected by M_C_. vitis and Z. vitis virophage after 12h pi.** 30 nm slices were acquired on a Scanning Electron Microscope FEI Teneo VS in « Serial Block-Face » mode (263 images 10nmx10nmx10nm and 3 energies deconvolution) and analysed using the VolumeScope™ software. The punctured VF (upper middle) surrounded by maturing M_C_. vitis virions as well as the cell nucleus (bottom right) are easily recognisable. The video can be found under the following link: https://mycore.core-cloud.net/index.php/s/juqBi1cvPfNF2UX

Table S1: Transpoviron proteins identified in the proteomes of host viruses and virophages

### Data Reporting

The annotated genomic sequences determined for this work have been deposited in the Genbank/EMBL/DDBJ database under the following accession numbers: M. vitis: MG807319, M. australiensis: MG807320, M. maliensis: MK978772, Z. vitis: MG807318, mv_C_tv: MG807316, ma_B_tv, MG807317.

The mass spectrometry proteomics data have been deposited to the ProteomeXchange Consortium via the PRIDE partner repository with the dataset identifier PXD009037(66).

Viral strains are available upon request.

## Acknowledgments

We thank Laurence De Marchi and Daouda Traore for providing the soil samples and Olivier Poirot for helpful discussions. We thank Dr. Elsa Garcin for reading and improving the manuscript. We also thank Dr. A. Kosta at the IMM imagery platform and F. Richard and A. Aouane for their expert assistance on the IBDM imaging platform and Dr. N. Brouilly for data acquisition that allowed producing the Supplementary movie. We also acknowledge the support of the bottom-up platform and informatics group of EDyP.

